## Supplemental Figures and Tables. for "Interplay between charge distribution and DNA in shaping HP1 paralog phase separation and localization"

**This PDF file includes:**

Figs. S1 to S5
Tables S1 to S2
Movies S1 to S6

### Supporting Information Figures

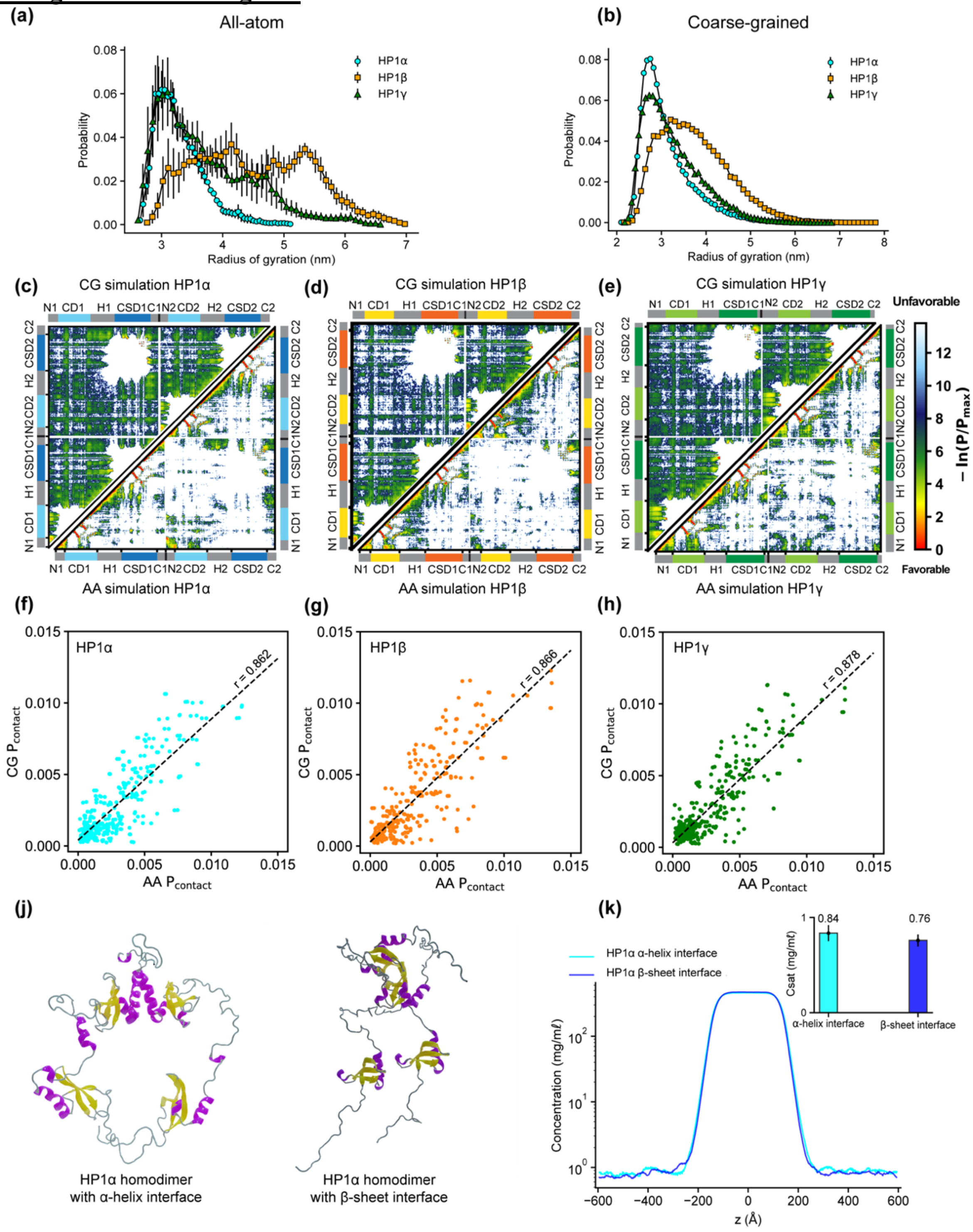


**Figure S1**: **(a, b)** $R_g$ distributions of HP1 paralogs in all-atom (AA) and coarse-grained (CG) simulations. **(c, d, f)** Contact maps for HP1 paralogs from all-atom simulations (bottom triangle) and coarse-grained simulations (top triangle). **(f, g, h)** Correlation of contact probability per residue between AA and CG simulations of the three HP1 paralogs. Pearson correlation coefficient (r) is shown above the regression line. Figs. S1a-h present the results from AA and CG simulations under dilute conditions. **(j)** Full-length HP1α homodimers in which the CSD-CSD domains dimerize via α-helix or β-sheet binding interfaces. **(h)** Density profiles of the two HP1α configurations in the CG phase coexistence simulations, conducted using the HPS-Urry model at 320 K and 100 mM salt concentration. The inset shows the saturation concentrations.

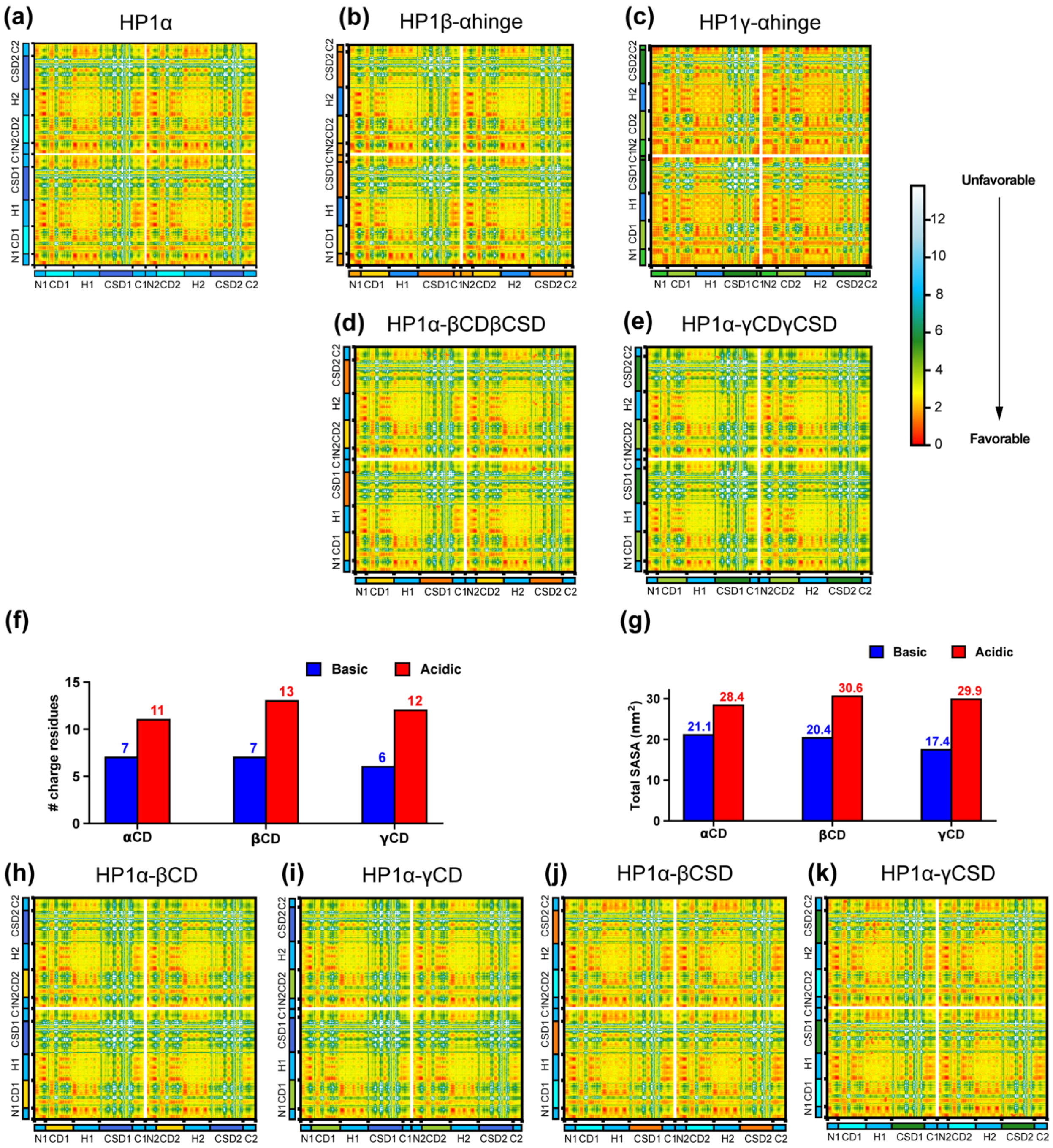


**Figure S2**: Intermolecular contact maps within the condensed phase of **(a)** wild-type HP1α, **(b, c)** chimeras in which the HP1α hinge was swapped with the hinge of HP1β or HP1γ, and **(d, e)** HP1α chimeras whose folded domains were replaced with those from either HP1β or HP1γ. **(e, f)** The number of basic and acidic residues and the total solvent-accessible surface areas (SASA) of basic and acidic residue on the CD domain of HP1 paralogs. **(h, i, j, k)** Intermolecular contact maps of HP1α constructs whose CD or CSD was replaced with the corresponding folded domain from either HP1β (HP1α-βCD and HP1α-βCSD, respectively) or HP1γ (HP1α-γCD and HP1α-γCSD, respectively).

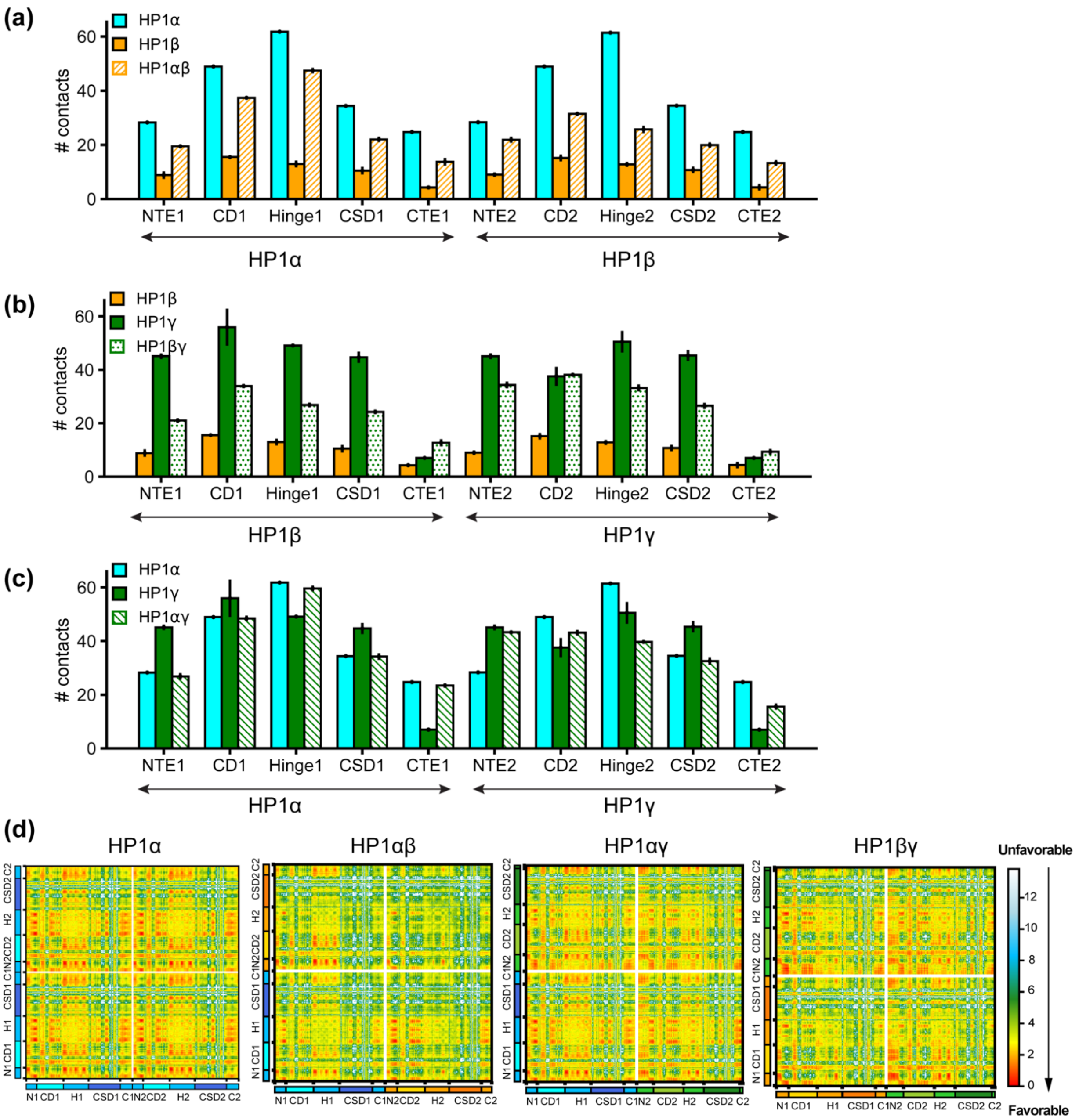


**Figure S3: (a, b, c)** Intermolecular contacts of different domains within condensates comprised of HP1αβ heterodimers (compared to HPα and HP1β homodimers), HP1αγ heterodimers (compared to HPα and HP1γ homodimers), and HP1βγ heterodimers (compared to HPβ and HP1γ homodimers). The labels under each plot mark the regions of the monomers in the heterodimer. **(d)** 2D intermolecular contact maps within condensates comprised of HP1α homodimers or HP1αβ, HP1αγ, and HP1βγ heterodimers.

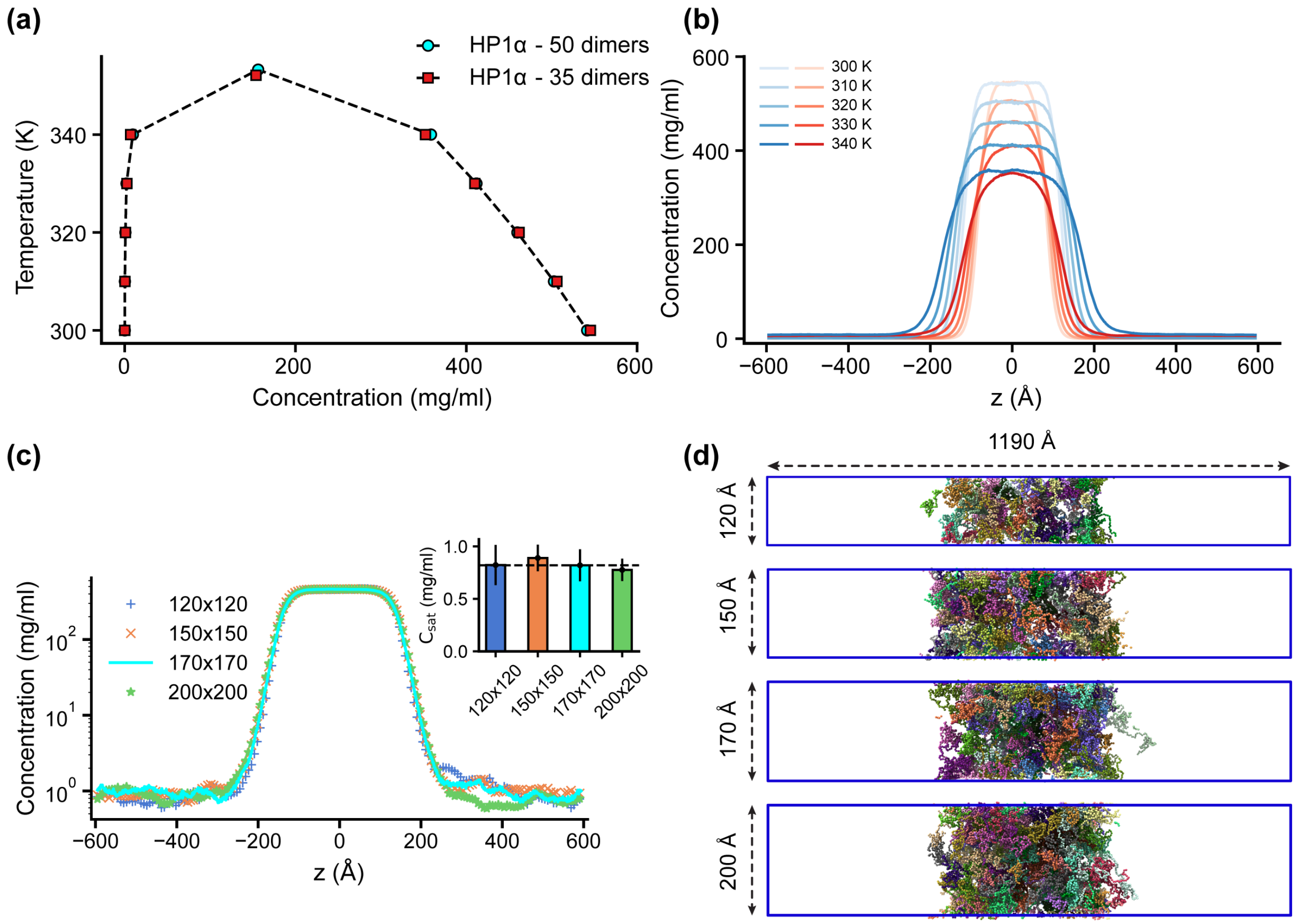


**Figure S4:** Finite-size analysis. **(a)** Phase diagrams for the HP1α homodimer (50 dimers) and for a system reduced in size by 30% (35 dimers), with the critical temperatures of 353.3 K and 352.1 K, respectively. **(b)** Density profiles of HP1α and its reduced size counterpart at various temperatures (blue – 50 dimers, red – 35 dimers). **(c, d)** Density profiles and snapshots of HP1α homodimer simulation with box dimensions of 170x170x1190 Å$^3$ and for the systems with the z-direction length fixed at 1190 Å and varying cross-sectional areas: 120x120, 150x150, and 200x200 Å$^2$. The black dashed line shows the simulated saturation concentration of wild-type HP1α homodimer in the box dimensions of 170x170x1190 Å$^3$. The simulations were conducted at 320 K and 100 mM salt concentrations. The error bars represent the standard deviation from triplicate simulation sets.

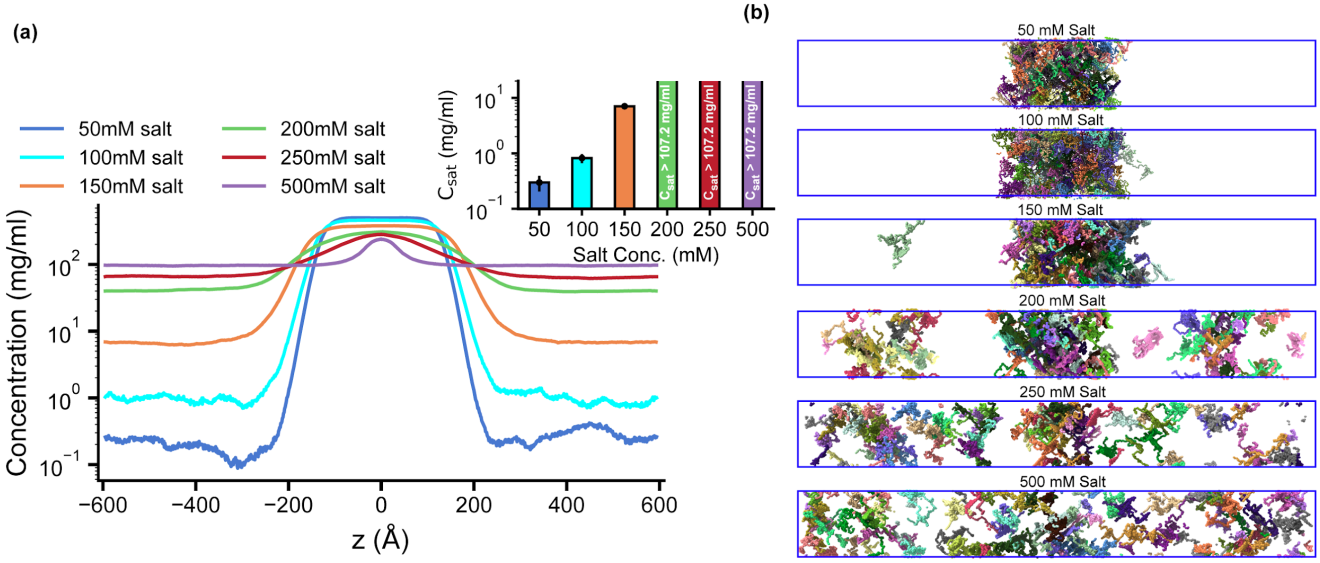


**Figure S5**: Salt-dependent effects on the LLPS of HP1α homodimers. **(a, b)** Density profiles and snapshots of HP1α homodimer simulations with box dimensions of 170x170x1190 Å$^3$ at differing salt concentrations, 50, 100, 150, 200, 250, and 500 mM, respectively. The simulations were conducted at 320 K using the HPS-Urry model. The error bars represent the standard deviation from triplicate simulation sets.

#### Supporting Information Tables

**Table S1**: Sequences of HP1 paralogs and their chimeras. The CD and CSD domains are highlighted in red and blue, respectively.

| | |
|---|---|
| HP1α (191aa) | MGKKTKRTADSSSSEDEEEYVVEKVLDRRVVKGQVEYLLKWKGFSEEHNTW EPEKNLDCPELISEFMKKYKKMKEGENNKPREKSESNKRKSNFSNSADDIK SKKKREQSNDIARGFERGLEPEKIIGATDSCGDLMFLMKWKDTDEADLVLA KEANVKCPQIVIAFYEERLTWHAYPEDAENKEKETAKS |
| HP1β (185aa) | MGKKQNKKKVEEVLEEEEEYVVEKVLDRRVVKGKVEYLLKWKGFSDEDNT WEPEENLDCPDLIAEFLQSQKTAHETDKSEGGKRKADSDSEDKGEESKPKK KKEESEKPRGFARGLEPERIIGATDSSGELMFLMKWKNSDEADLVPAKEAN VKCPQVVISFYEERLTWHSYPSEDDDKKDDKN |
| HP1γ (183aa) | MASNKTTLQKMGKKQNGKSKKVEEAEPEEFVVEKVLDRRVVNGKVEYFLKW KGFTDADNTWEPEENLDCPELIEAFLNSQKAGKEKDGTKRKSLSDSESDDS KSKKKRDAADKPRGFARGLDPERIIGATDSSGELMFLMKWKDSDEADLVLA KEANMKCPQIVIAFYEERLTWHSCPEDEAQ |
| HP1α-βIDRs (185aa) | MGKKQNKKKVEEVLEEEEEYVVEKVLDRRVVKGQVEYLLKWKGFSEEHNT WEPEKNLDCPELISEFMQSQKTAHETDKSEGGKRKADSDSEDKGEESKPKK KKEESEKPRGFERGLEPEKIIGATDSCGDLMFLMKWKDTDEADLVLAKEAN VKCPQIVIAFYEERLTWHAYPSEDDDKKDDKN |
| HP1α-γIDRs (179aa) | MASNKTTLQKMGKKQNGKSKKVEEAEPEEYVVEKVLDRRVVKGQVEYLLKW KGFSEEHNTWEPEKNLDCPELISEFMKAGKEKDGTKRKSLSDSESDDSKSK KKRDAADKPRGFERGLEPEKIIGATDSCGDLMFLMKWKDTDEADLVLAKEA NVKCPQIVIAFYEERLTWHAYPEDEAQ |
| HP1α-βHinge (186aa) | MGKKTKRTADSSSSEDEEEYVVEKVLDRRVVKGQVEYLLKWKGFSEEHNTW EPEKNLDCPELISEFMQSQKTAHETDKSEGGKRKADSDSEDKGEESKPKKK KEESEKPRGFERGLEPEKIIGATDSCGDLMFLMKWKDTDEADLVLAKEANV KCPQIVIAFYEERLTWHAYPEDAENKEKETAKS |
| HP1α-γHinge (178aa) | MGKKTKRTADSSSSEDEEEYVVEKVLDRRVVKGQVEYLLKWKGFSEEHNTW EPEKNLDCPELISEFMKAGKEKDGTKRKSLSDSESDDSKSKKKRDAADKPR GFERGLEPEKIIGATDSCGDLMFLMKWKDTDEADLVLAKEANVKCPQIVIA FYEERLTWHAYPEDAENKEKETAKS |
| HP1β-αHinge (190aa) | MGKKQNKKKVEEVLEEEEEYVVEKVLDRRVVKGKVEYLLKWKGFSDEDNT WEPEENLDCPDLIAEFLKKYKKMKEGENNKPREKSESNKRKSNFSNSADDI KSKKKREQSNDIARGFARGLEPERIIGATDSSGELMFLMKWKNSDEADLVP AKEANVKCPQVVISFYEERLTWHSYPSEDDDKKDDKN |
| HP1γ-αHinge (196aa) | MASNKTTLQKMGKKQNGKSKKVEEAEPEEFVVEKVLDRRVVNGKVEYFLKW KGFTDADNTWEPEENLDCPELIEAFLNSQKKYKKMKEGENNKPREKSESNK RKSNFSNSADDIKSKKKREQSNDIARGFARGLDPERIIGATDSSGELMFLM KWKDSDEADLVLAKEANMKCPQIVIAFYEERLTWHSCPEDEAQ |

| | |
|---|---|
| HP1α-βCDβCSD (190aa) | MGKKTKRTADSSSSEDEEEYVVEKVLDRRVVKGKVEYLLKWKGFSDEDNTW EPEENLDCPDLIAEFLKKYKKMKEGENNKPREKSESNKRKSNFSNSADDIK SKKKREQSNDIARGFARGLEPERIIGATDSSGELMFLMKWKNSDEADLVPA KEANVKCPQVVISFYEERLTWHSYPEDAENKEKETAKS |
| HP1α-γCDγCSD (193aa) | MGKKTKRTADSSSSEDEEEFVVEKVLDRRVVNGKVEYFLKWKGFTDADNTW EPEENLDCPELIEAFLNSQKKYKKMKEGENNKPREKSESNKRKSNFSNSAD DIKSKKKREQSNDIARGFARGLDPERIIGATDSSGELMFLMKWKDSDEADL VLAKEANMKCPQIVIAFYEERLTWHSCPEDAENKEKETAKS |
| HP1α-CDβ (190aa) | MGKKTKRTADSSSSEDEEEYVVEKVLDRRVVKGKVEYLLKWKGFSDEDNTW EPEENLDCPDLIAEFLKKYKKMKEGENNKPREKSESNKRKSNFSNSADDIK SKKKREQSNDIARGFERGLEPEKIIGATDSCGDLMFLMKWKDTDEADLVLA KEANVKCPQIVIAFYEERLTWHAYPEDAENKEKETAKS |
| HP1α-CDγ (193aa) | MGKKTKRTADSSSSEDEEEFVVEKVLDRRVVNGKVEYFLKWKGFTDADNTW EPEENLDCPELIEAFLNSQKKYKKMKEGENNKPREKSESNKRKSNFSNSAD DIKSKKKREQSNDIARGFERGLEPEKIIGATDSCGDLMFLMKWKDTDEADL VLAKEANVKCPQIVIAFYEERLTWHAYPEDAENKEKETAKS |
| HP1α-CSDβ (191aa) | MGKKTKRTADSSSSEDEEEYVVEKVLDRRVVKGQVEYLLKWKGFSEEHNTW EPEKNLDCPELISEFMKKYKKMKEGENNKPREKSESNKRKSNFSNSADDIK SKKKREQSNDIARGFARGLEPERIIGATDSSGELMFLMKWKNSDEADLVPA KEANVKCPQVVISFYEERLTWHSYPEDAENKEKETAKS |
| HP1α-CSDγ (191aa) | MGKKTKRTADSSSSEDEEEYVVEKVLDRRVVKGQVEYLLKWKGFSEEHNTW EPEKNLDCPELISEFMKKYKKMKEGENNKPREKSESNKRKSNFSNSADDIK SKKKREQSNDIARGFARGLDPERIIGATDSSGELMFLMKWKDSDEADLVLA KEANMKCPQIVIAFYEERLTWHSCPEDAENKEKETAKS |

**Table S2**: Double-stranded DNA sequence used in the simulations.

| | |
|---|---|
| 147 bp 601 dsDNA | CTGGAGAATCCCGGTGCCGAGGCCGCTCAATTGGTCGTAGACAGCTCTAGC ACCGCTTAAACGCACGTACGCGCTGTCCCCCGCGTTTTAACCGCCAAGGGG ATTACTCCCTAGTCTCCAGGCACGTGTCAGATATATACATCCTGT |

#### Supporting Information Movies

**Movie S1.** CG simulations of HP1 paralogs. The CSD-CSD domains were aligned and fixed for visualization. The folded and disordered domains are shown in surface and tube representations.
**Movie S2**. CG coexistence phase simulations of HP1α, HP1β, and HP1γ paralogs.
**Movie S3**. CG coexistence phase simulations of HP1α, HP1β, and HP1γ paralogs with 147 bp dsDNA.
**Movie S4**. CG coexistence phase simulations and density profiles of HP1α (cyan) and HP1β (orange) paralogs on the left and HP1α (cyan) and HP1γ (green) paralogs on the right at an equimolar concentration.
**Movie S5**. CG coexistence phase simulations and density profiles of HP1α (cyan) and HP1β (orange) paralogs on the left and HP1α (cyan) and HP1γ (green) paralogs on the right at an equimolar concentration with 147 bp dsDNA.
**Movie S6**. CG coexistence phase simulations and density profiles of HP1α (cyan) and HP1β (orange), and HP1γ (green) paralogs at an equimolar concentration in the absence (left) and presence (right) of 147 bp dsDNA (red).
